# *Rnf149-related* is an FGF/MAPK-independent regulator of pharyngeal muscle fate specification

**DOI:** 10.1101/2022.01.07.475354

**Authors:** Burcu Vitrinel, Christine Vogel, Lionel Christiaen

**Author notes:** equally contributing authors (Christine Vogel, Lionel Christiaen).

## Abstract

During embryonic development, cell fate specification gives rise to dedicated lineages that underlie tissue formation. In olfactores, which comprise tunicates and vertebrates, the cardiopharyngeal field is formed by multipotent progenitors to both cardiac and branchiomeric muscles. The ascidian *Ciona* is a powerful model to study the cardiopharyngeal fate specification with cellular resolution, as only 2 pairs of cardiopharyngeal multipotent progenitors give rise to the heart and to pharyngeal muscles (*aka* atrial siphon muscles, ASM). These progenitors are multilineage primed, in as much as they express a combination of early ASM- and heart-specific transcripts that become restricted to their corresponding precursors, following oriented asymmetric divisions. Here, we identify the primed gene *Rnf149-related* (*Rnf149-r*), which becomes restricted to the heart progenitors, but appears to regulate pharyngeal muscle fate specification in the cardiopharyngeal lineage. CRISPR/Cas9-mediated loss knock-out of *Rnf149-r* function impairs atrial siphon muscle morphogenesis, and down-regulates *Tbx1/10* and *Ebf*, two key determinants of the pharyngeal muscle fate, while upregulating heart-specific gene expression. These phenotypes are reminiscent of loss of FGF-MAPK signaling in the cardiopharyngeal lineage, and integrated analysis of lineage-specific bulk RNA-seq profiling of loss-of-function perturbations identified a significant overlap between FGF-MAPK and Rnf149-r targets. However, functional interaction assays suggested the Rnf149-r does not directly modulate the activity of the FGF-MAPK-Ets1/2 pathway. Instead, we propose that *Rnf149-r* acts both in parallel to the FGF-MAPK signaling on shared targets, as well as on FGF-MAPK-independent targets through (a) separate pathway(s).

## Introduction

During vertebrate development, the heart arises from distinct first and second heart fields (Buckingham, 2016; Kelly, 2020; Kelly et al., 2014). Clonal analyses have shown that first and second heart field progenitor cells arise from independent pools of multipotent *Mesp1*+ progenitors (Devine et al., 2014; Lescroart et al., 2014). However, common pools of progenitors give rise to the second heart field and the branchiomeric/pharyngeal muscles (Lescroart et al., 2010; Lescroart et al., 2015), referred to as cardiopharyngeal lineages.

Here, we leverage the simplicity of the cardiogenic lineage of *Ciona*, a simple tunicate and the closest living relatives to vertebrates among chordates (Delsuc et al., 2006; Putnam et al., 2008), to study cardiopharyngeal cell fate choices. *Ciona* allows us to study conserved early stages of cardiopharyngeal development with exceptional spatial and temporal resolution, and therefore emerged as a suitable model organism to understand developmental fate choices between cardiac and pharyngeal muscle cells (Kaplan et al., 2015). In *Ciona*, early lineage commitment typically restricts the competence of progenitors prior to lineage amplification by proliferation; by contrast, multipotent progenitors are amplified prior to fate specification in mammalian embryos (Evans Anderson and Christiaen, 2016). Like their vertebrate counterparts, *Ciona’s* cardiopharyngeal lineage stems from multipotent progenitors, the trunk ventral cells (TVCs), which emerge from *Mesp*+ mesodermal progenitors. The TVCs are induced by Fibroblast growth factor/Mitogen-activated protein kinase (FGF/MAPK) signaling and migrate as bilateral pairs of cells, until the left and right pairs meet at the ventral midline and resume cell divisions (**Figure 1a**) (Christiaen et al., 2008; Davidson et al., 2005; Satou et al., 2004; Stolfi et al., 2010). Late TVCs undergo oriented asymmetric divisions that produce first heart precursors (FHPs) and secondary TVCs (STVCs), which then give rise to second heart precursors (SHPs) and pharyngeal muscle precursors (aka atrial muscle founder cells, ASMFs) **(Figure 1a)**. STVCs activate *Tbx1/10*, the homolog of human *TBX1* that is required for cell-specific expression of the pharyngeal muscle determinant *Ebf* expression (Razy-Krajka et al., 2014; Stolfi et al., 2010; Wang et al., 2013). The FGF/MAPK signaling pathway is a key regulator of successive cardiopharyngeal fate decisions in *Ciona*. FGF/MAPK signaling is required for the induction of first and second-generation multipotent cardiopharyngeal progenitors (aka TVCs, and STVCs), and for the subsequent activation of the siphon/pharyngeal muscle program in their corresponding progenitors (Davidson et al., 2006; Razy-Krajka et al., 2018).

**Figure 1:**
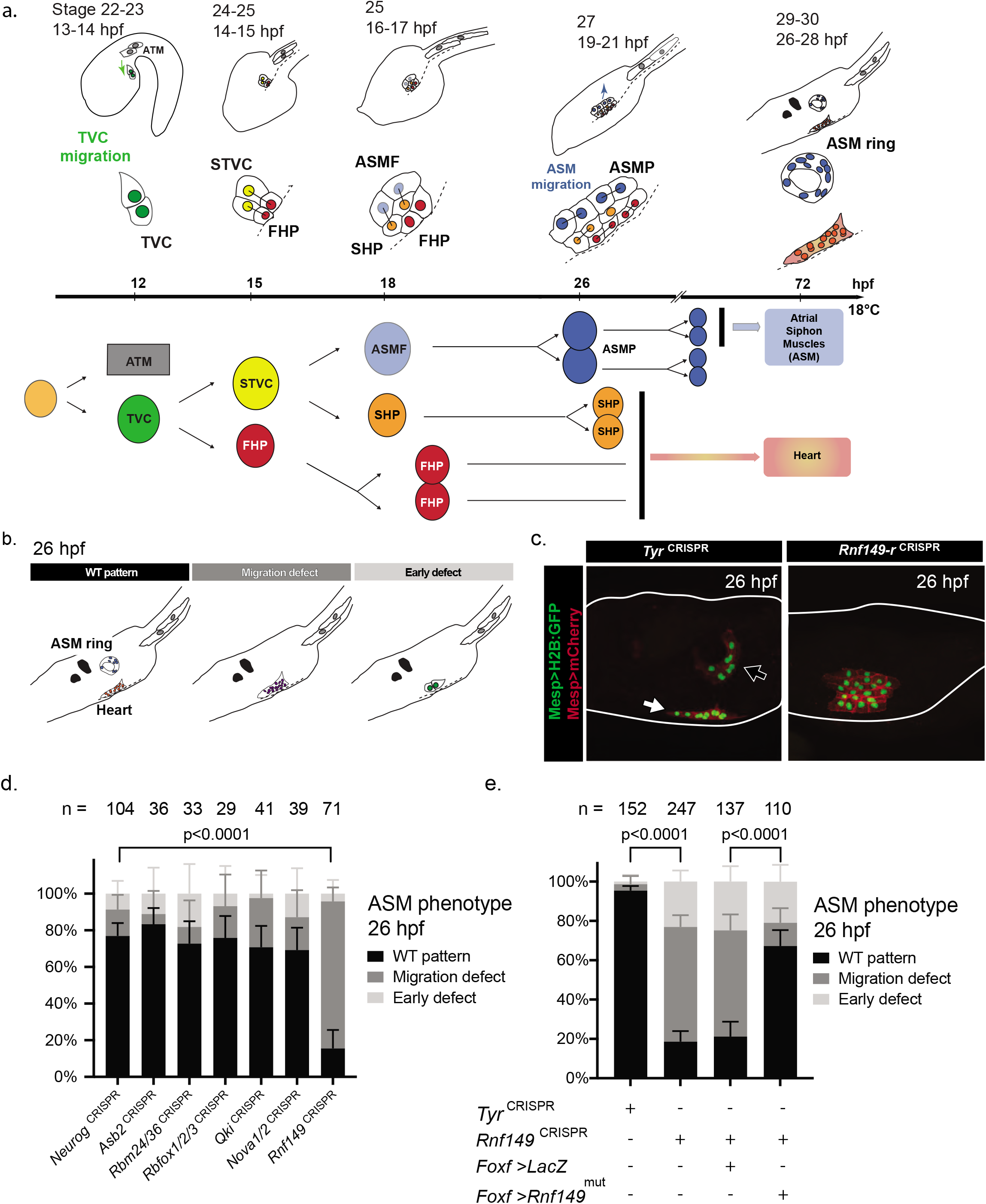
*Rnf149-r*^CRISPR^ causes ASM migration defects. a) Schematic of *Ciona* development showing asymmetric cell divisions and resulting cell fates of the cardiopharyngeal mesoderm (CPM). Stages are set according to Hotta et al. (2007) with hours post fertilization (hpf) at 18°C (**Supp table 2**). Anterior tail muscle (ATM, gray), trunk ventral cell (TVC, green), secondary TVC (STVC, yellow), first heart precursor (FHP, red), second heart precursor (SHP, orange), atrial siphon precursor cells (ASMF, blue). Dashed lines indicate the ventral midline. b) Schematic representation of phenotypes scored in Figure 1c. c) Tyr^CRISPR^ used as control. Cardiopharyngeal lineage are marked by mCherry and GFP driven by *Mesp*. H2B::GFP (green) and hCD4::mCherry (red) accumulate in the nuclei and at the cell membrane, respectively. Arrow indicates the heart progenitors and the open arrow indicates the ASM ring. d) Histogram with phenotype proportions. *Neuro*g^CRISPR^ are used as control, a gene that is known to be inactive in the cardiopharyngeal lineage. First 5 genes scored compared to *Neurog*^CRISPR^ are not significant per Fisher exact test while *Rnf149-r*^CRISPR^ is significant with *p*-value<0.0001. Experiments are performed in biological replicates and “n=” represents the total numbers of individual halves scored per condition. Error bars represent 95% Wilson method of confidence interval for proportions. e) Corresponding histogram with phenotype proportions. Experiments are performed in biological replicates. “n=” represents the total numbers of individual halves scored per condition. Error bars represent 95% Wilson method of confidence interval for proportions.

Over the past decade, we have extensively documented gene expression dynamics, and begun to decipher the underlying gene regulatory networks that govern early cardiopharyngeal development in *Ciona*. A key feature of the transcriptome dynamics that determine cardiopharyngeal transitions is multilineage priming, whereby multipotent cardiopharyngeal progenitors co-express early key regulators of the cardiac- and pharyngeal muscle-specific programs (Razy-Krajka et al., 2014; Wang et al., 2019). We surmise that multilineage transcriptional priming, while contributing to multipotency, also poses a challenge for subsequent fate specification following cell divisions, as fate-restricted progenitors inherit gene products that belong to the alternative fates, and might interfere commitment to a cardiac or pharyngeal muscle identity. For instance, single cell RNA-seq datasets indicated that first and second heart precursors inherit pharyngeal muscle-specific mRNAs that were down-regulated with varying dynamics after cell division, and upon commitment to a cardiac identity (Wang et al., 2019). We thus hypothesize that cell-type specific post-transcriptional regulatory mechanisms contribute to early cardiopharyngeal development, by remodeling inherited transcriptomes and proteomes upon fate specification and commitment.

Here, we focused on candidate transcriptional regulators showing differential gene expression in the cardiopharyngeal lineage, and identified *Rnf149-related* (hereafter referred to as *Rnf149-r*), a previously uncharacterized gene, as a necessary determinant of pharyngeal muscle identity. *Rnf149-r* is a transcriptionally primed heart marker in the cardiopharyngeal lineage (Wang et al., 2019) also encodes the postplasmic RNA (aka posterior end mark, PEM) *Pen-1* (Yamada et al., 2005). Finally, *Rnf149-r/Pen-1* is also dynamically expressed in various tissues during embryogenesis, including the central nervous system, the notochord and the epidermis, further suggesting pleiotropic functions. Rnf149-r’s predicted protein structure revealed an atypical RNF organization, with a protease-associated domain but lacking the catalytic RING domain. In the cardiopharyngeal lineage, CRISPR/Cas9-mediated loss of *Rnf149-r* function disrupted pharyngeal muscle specification, most likely through inhibition of *Ebf* gene expression. The effects of *Rnf149-r^CRISPR^* partially phenocopied loss of FGF/MAPK signaling, including a significant overlap of dysregulated genes from bulk RNA-seq experiments on FACS-purified cells. Finally, functional interaction assays indicated that *Rnf149-r* acts in parallel to the FGFR-MEK-Ets1/2 pathway upstream of *Ebf* activation, thus revealing the existence of FGF/MAPK-independent regulatory inputs into pharyngeal muscle specification.

## Results

### CRISPR/Cas9-mediated mutagenesis identifies the pharyngeal muscle determinant Rnf149-r

We previously observed extensive multilineage transcriptional priming in multipotent cardiopharyngeal progenitors (Wang et al., 2019). This led us to hypothesize that lineage-specific post-transcriptional regulatory mechanisms contribute to remodelling transcriptomes and proteomes during heart vs. pharyngeal muscle fate decisions. To identify candidate post-transcriptional regulators, we catalogued genes encoding proteins annotated as RNA-binding and/or involved in ubiquitination pathways (GO terms GO:0003723 and GO:0016567, respectively), using the ANISEED database of GOSlim annotations curated using data from InterPro and UniProt (**Supp. table 1**) (Blum et al., 2021; Brozovic et al., 2016; UniProt Consortium, 2019). We integrated this table, containing 945 candidate genes, with previous cardiopharyngeal lineage-specific single cell RNA-seq (scRNA-seq) data, to identify candidate heart- and pharyngeal muscle-specific post-transcriptional regulators (Wang et al., 2019) (**Supp. tables 1, 2**). In a pilot approach, we selected 14 cardiopharyngeal genes that encode either ubiquitin ligase-related proteins (*Rnf149-r, Asb2, Bag3/4, Rbms1/2/3*) or RNA-binding proteins (*Nova, Rbfox1/2/3, Rbms1/2/3, Rbm24/38, Qki, Ube2ql1, Pcbp3, Ube2j1, Otud3, Psmd14*). We performed dual fluorescent *in situ* hybridization and immunohistochemistry (FISH-IHC) to verify their predicted expression in the heart and/or pharyngeal muscle precursors **(Figure S1)**. We conducted lineage-specific loss-of-function analyses using the CRISPR/Cas9 system to target 6 of 15 candidate regulators (Gandhi et al., 2017; Gandhi et al., 2018; Haeussler et al., 2016; Stolfi et al., 2014). We collected swimming larvae (Stage 29/30; 26 hours post-fertilization), and scored the morphology of the cardiopharyngeal lineage, notably the presence or absence of the conspicuous pharyngeal/atrial siphon muscle (ASM) rings **(Fig. 1b,c**; (Stolfi et al., 2010)). We targeted the neurogenic bHLH factor-coding gene *Neurogenin* (*Neurog*^CRISPR^) as a control, since it is neither expressed in the cardiopharyngeal mesoderm nor thought to be involved in cardiopharyngeal development (Gandhi et al., 2017).

Lineage-specific CRISPR/Cas9-mediated mutagenesis of *Rnf149-r* showed the most penetrant phenotype, characterized by disrupted pharyngeal muscle morphogenesis (Chi-square test, p-value < 0.0001; **Fig. 1d**). *Rnf149-r* mutagenesis caused pharyngeal muscle morphogenesis defects, whereby the cells failed to migrate toward the atrial siphon placode (ASP) and form the ASM rings or crescents observed in control larvae (Stolfi et al., 2010). ASM rings were present in control conditions targeting either *Neurog* or *Tyrosinase* (*Tyr*), which is also inactive in the cardiopharyngeal lineage (**Fig. 1b-e**). To confirm the specificity of the *Rnf149-r^CRISPR^* phenotype, we targeted *Rnf149-r* using 2 sgRNAs targeting different positions of the coding sequence (**Figure S2a**). These sgRNAs produced similar pharyngeal muscle morphogenesis defects, whether used in combination or separately, indicating that both reagents contribute to the phenotype and are specific to *Rnf149-r* (**Figure S2b**). To further ascertain specificity, we expressed the CRISPR/Cas9 reagents alongside a rescue construct, consisting of a CRISPR/Cas9-resistant form of an *Rnf149-r* cDNA, with mutations in the protospacer adjacent motifs (PAMs) of both sgRNAs used (*Rnf149-r*^mut^), and expressed under the cardiopharyngeal progenitor-specific *Foxf* enhancer (Beh et al., 2007). Remarkably, the proportion of larvae showing signs of normal ASM morphogenesis increased from ~20% in *Rnf149-r^CRISPR^* animals to ~70% following co-expression of *Rnf149-r*^mut^ (**Fig. 1e**). This experiment demonstrated that the observed pharyngeal muscle phenotype is specifically caused by the loss of *Rnf149-r* function in the cardiopharyngeal lineage.

Consistent with a potential role in cardiopharyngeal development, the uncharacterized gene *Ring finger protein 149 related* (*Rnf149-r*) is transcriptionally primed in multipotent progenitors of tailbud embryos (stage 22), and restricted to the heart progenitors in swimming larvae (**Figure S3**) (Wang et al., 2019). To understand whether the phenotype is primarily caused by late migration, or early fate specification defects, we assayed expression of the essential ASM determinant and specific marker *Ebf*, in hatching larvae (stage 26; (Racioppi et al., 2019; Razy-Krajka et al., 2014; Stolfi et al., 2010)). The ASM-specific factor *Ebf* is necessary and sufficient to suppress the heart program, and impose the pharyngeal muscle fate in the cardiopharyngeal lineage (Razy-Krajka et al., 2014; Stolfi et al., 2010). *Rnf149-r* mutagenesis caused lineage-specific loss of *Ebf* expression, which typically produced ectopic cardiac specification and ASM fate specification defects, thus abolishing migration toward the ASP (**Figure 2a, b**) (Razy-Krajka et al., 2014; Stolfi et al., 2014).

**Figure 2:**
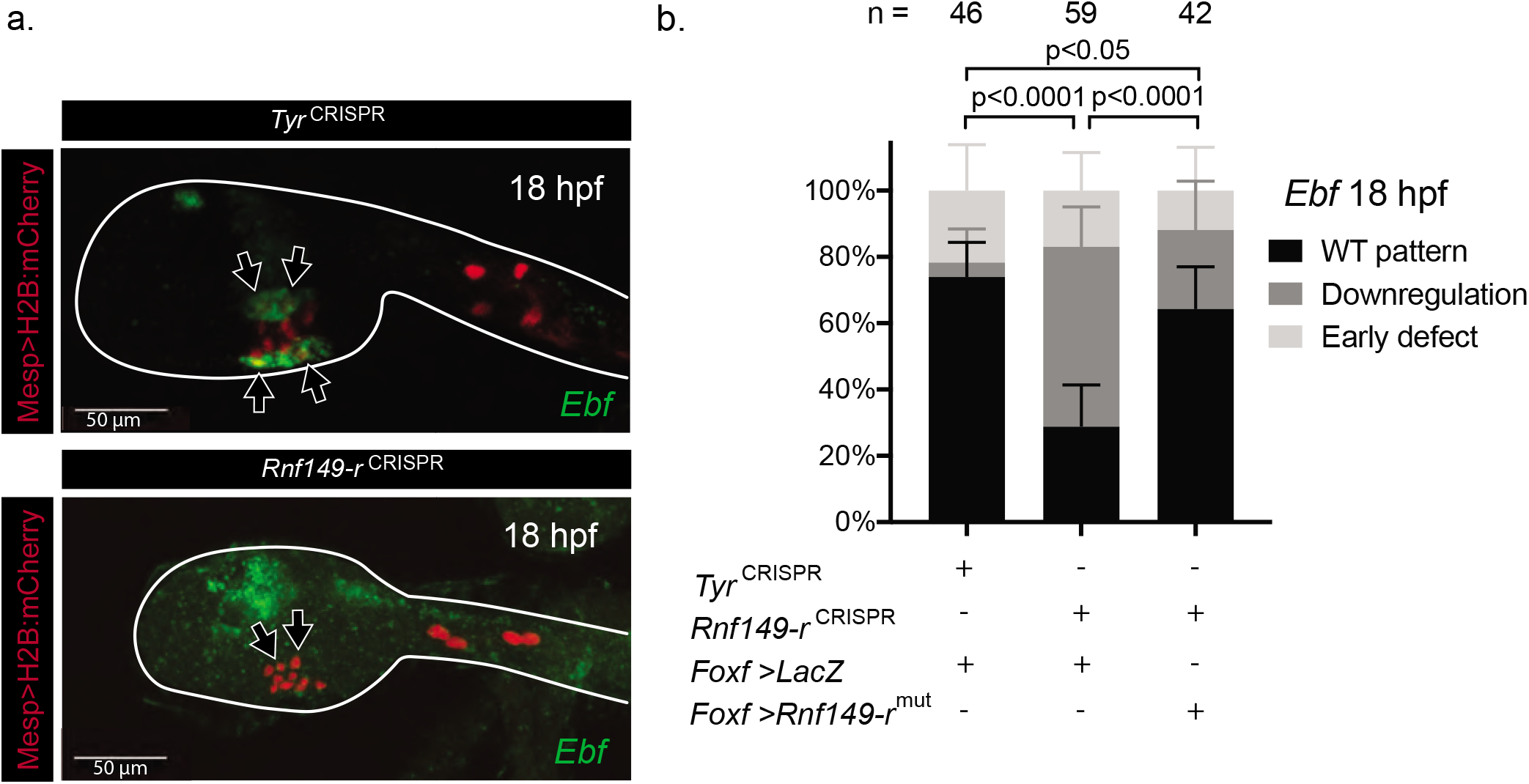
*Rnf149-r*^CRSPR^ causes behavioral defects in the cardiopharyngeal lineage. a) In situ hybridization using an *Ebf* probe in green. *Tyr*^CRISPR^ used as control. Cardiopharyngeal lineage are marked by mCherry driven by *Mesp* and revealed by mCherry antibody in red. H2B::mCherry accumulates in the nuclei. Scale bar is 50 μm. Arrows mark ASMFs. b) Corresponding histogram with phenotype proportions. “n=” represents the total numbers of individual halves scored per condition. Error bars represent 95% Wilson method of confidence interval for proportions.

As we observed phenotypic defects at 18 and 26 hpf, we asked whether earlier cardiopharyngeal development is affected by loss of *Rnf149-r* function. We tested TVC migration and expression of the TVC marker *Hand-r* expression at the late tailbud stage (12 hpf at 18°C). However, we did not observe any difference between the experimental and control animals (**Figure S4**). In light of these results, we propose that *Rnf149-r* function is necessary for the transition to the pharyngeal muscle fate from a multipotent cardiopharyngeal progenitor state.

### Rnf149-r encodes an atypical ubiquitin ligase-related protein

We identified *Rnf149-r* for CRISPR/Cas9 mutagenesis as a candidate post-transcriptional regulator, because it was annotated as a RING-finger domain containing protein, which typically comprise E3-ubiquitin ligases. However, upon closer inspection, we found that the predicted *Ciona* Rnf149-r protein lacks a RING domain, but contains a Protease-Associated (PA) domain (**Figure S5**). PA domains in humans and other higher vertebrates can co-exist with RING domains, as well as other functionally active domains, such as EGF, RZF family and transferrin receptor (Mahon and Bateman, 2000).

Like other organisms, the *Ciona* genome encodes a variety of PA-domain-containing proteins. This domain in *Ciona* occurs as the only defined domain in 2 predicted proteins, including Rnf149-r. The other PA-domain proteins also contain associated glycosidase domains, Zn-independent exopeptidase domains, transferrin receptor-like dimerization domains and/or RING domains (**Figure S5, Supp. table 3**). Our sequence analyses showed that Rnf149-r has one homolog with similar domain architecture in Ciona, Rnf150, prompting us to hypothesize that Rnf149-r may act as a natural dominant-negative inhibitor of Rnf150 function. However, CRISPR/Cas9-mediated loss of Rnf150 function did not cause any overt phenotype, nor did it rescue loss of Rnf149-r function, leading us to rule out Rnf150 as mediator of the Rnf149-r^CRISPR^ phenotype (**Figure S6**).

### Rnf149-r regulates cardiopharyngeal fates independently of FGF/MAPK signaling

Fibroblast growth factor/Mitogen-activated protein kinase (FGF/MAPK) signaling is a key regulator of cardiopharyngeal fates in *Ciona*, with established roles in early *Mesp*+ mesoderm specification, and multipotent progenitor induction and migration (Christiaen et al., 2008; Davidson et al., 2006). Sustained FGF-MAPK activity leads to localized *Ebf* expression in ASM precursors, while its exclusion from first and second heart precursors permits cardiac specification (Razy-Krajka et al., 2018; Wang et al., 2019). MAPK activity in early pharyngeal muscle progenitors initiates *Ebf* expression, until Ebf accumulation permits MAPK-independent auto-activation. This switch is surmised to explain the transition from the multipotent state to committed pharyngeal muscle fate (Razy-Krajka et al., 2018).

The *Rnf149-r^CRISPR^* phenotype resembles loss of MAPK function, as observed following lineage-specific misexpression of a dominant-negative form of the FGF receptor, or by treatment with the MEK inhibitor U0126 (Razy-Krajka et al., 2018). Moreover, *in vitro* studies showed that human RNF149 interacts with and induces ubiquitination of the classic regulator of Mek1/2 and MAPK signaling, *Braf* (Hong et al., 2012). We thus hypothesized that *Rnf149-r* regulates the pharyngeal muscle fate choice by interacting with FGF/MAPK signaling.

To test these hypotheses, we overexpressed constitutively active forms of M-Ras and Mek1/2 in parallel with *Rnf149-r*^CRISPR^, and used *Ebf* expression as the readout of pharyngeal muscle fate specification. Overexpression of constitutively active forms of either M-Ras or Mek1/2 suffices to cause ectopic *Ebf* expression in the cardiopharyngeal lineage, and abolish the heart fate (Razy-Krajka et al., 2018) (**Figure 3**). We first combined *Rnf149-r*^CRISPR^ with overexpression of a defined constitutively active form of M-Ras, M-Ras^G22V^ (called M-Ras^CA^ here on), which is the only Ras homolog in *Ciona* and acts in the FGF/MAPK pathway (Keduka et al., 2009). We also overexpressed a constitutively active form of Mek1/2, Mek1/2^S220E,S216D^ (Mek^CA^ here on) (Razy-Krajka et al., 2018), a key regulator of MAPK activity downstream of M-Ras (**Figure 3a**). We expressed these constructs using the TVC-specific *Foxf* enhancer to restrict the misexpression of the constitutively active mutants to the TVCs and their progeny. Accordingly, we did not observe any unrelated, early cardiopharyngeal development defects. In either case, the effects of *Rnf149-r*^CRISPR^ dominated ectopic activation of the Ras-Mek pathway and blocked *Ebf* expression (**Figure 3**). The dominance of the Rnf149^CRISPR^ phenotype was even more clearly observable when using *Foxf>LacZ* to label transfected cells and account for mosaicism (**Figure S7**). These results indicate that Rnf149-r function is required either in parallel to the FGF/MAPK pathway, or downstream of Mek, for proper *Ebf* expression, and by extension for pharyngeal muscle specification.

**Figure 3:**
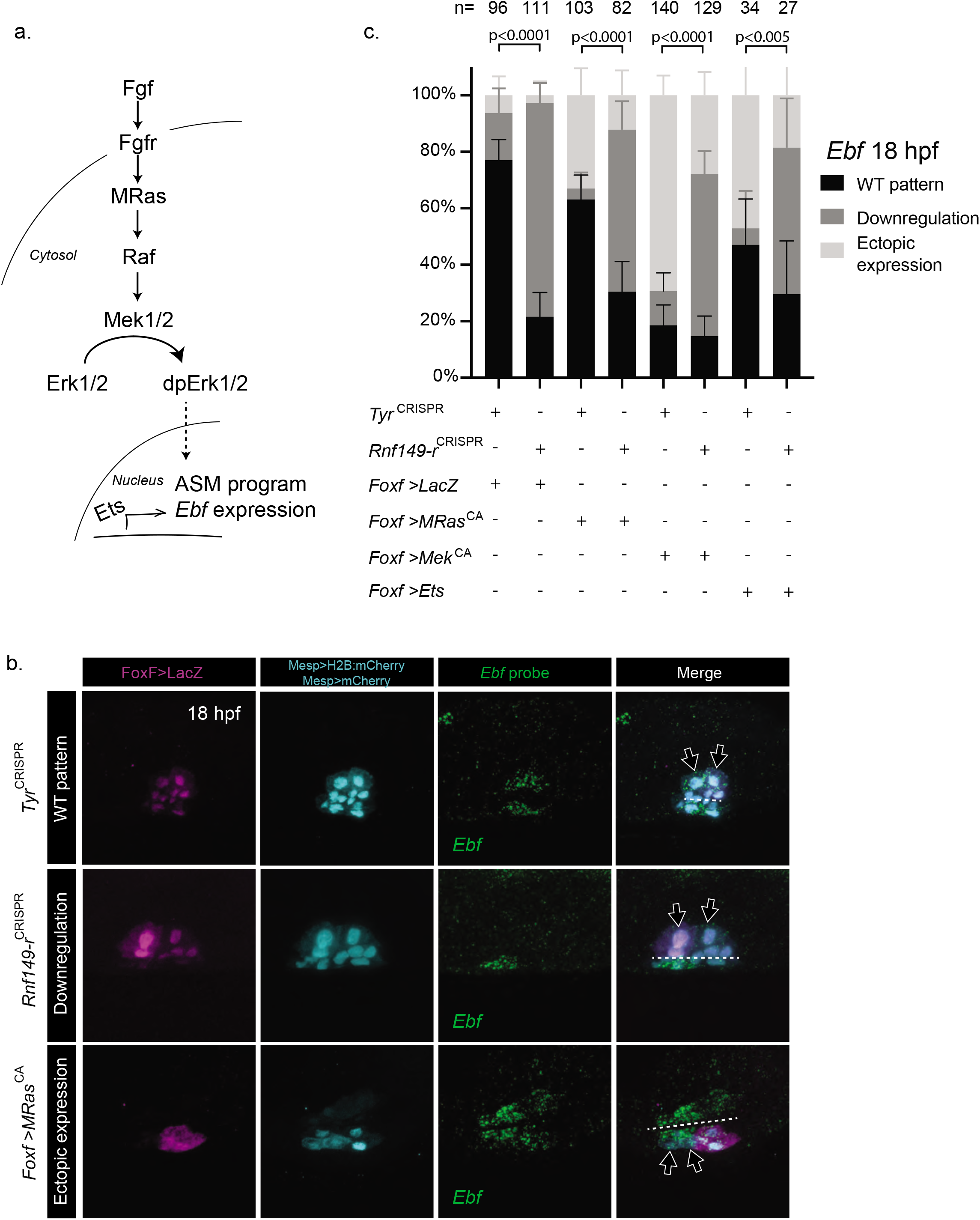
*Rnf149-r* acts in parallel to the FGF/MAPK pathway. a) Schematic representation of the FGF/MAPK pathway. b) In situ hybridization using an *Ebf* probe. *Tyr*^CRISPR^ are used as control. Cardiopharyngeal lineage is marked by mCherry and LacZ driven by *Mesp* and *FoxF*, revealed by mCherry and Beta-gal antibodies in magenta and cyan, respectively. H2B::mCherry and hCD4::mCherry (cyan) accumulate in the nuclei and at the cell membrane, respectively. Dashed lines indicate the ventral midline. Arrows mark ASMFs. c) Corresponding histogram with phenotype proportions. “n=” represents the total numbers of individual halves scored per condition. Error bars represent 95% Wilson method of confidence interval for proportions.

The transcription factor *Ets1/2* is a known downstream effector of the FGF/MAPK pathway, presumed to control cardiopharyngeal development in *Ciona* (Davidson et al., 2006; Miya and Nishida, 2003). We tested possible functional interactions between *Rnf149-r* and *Ets1/2*, and found that *Rnf149-r*^CRISPR^ inhibits the ectopic *Ebf* expression phenotype obtained with *Ets1/2* overexpression, as well (**Figure 3**).

This systematic dominance of the *Rnf149-r^CRISPR^* phenotype over gain of either M-Ras, Mek1/2 or Ets1/2 function suggested that the uncharacterized protein Rnf149-r acts in parallel to the FGF/MAPK pathway upstream of *Ebf* activation during pharyngeal muscle specification. This is consistent with the above conclusion Rnf149-r functions later than late tailbud stage, since FGF/MAPK is already active and necessary for multipotent progenitor induction and maintenance (Davidson et al. 2006; Razy-Krajka et al. 2018).

### Rnf149-r regulates both MAPK-dependent and independent genes

To explore the broader transcriptional impact of *Rnf149-r* loss-of-function, we performed lineage-specific bulk RNA-seq experiment by FACS-purifying cardiopharyngeal cells following CRISPR/Cas9-mediated mutagenesis of either *Rnf149-r*, or *Tyrosinase* as a control, in biological triplicates. Out of 15,232 genes quantified, 190 were significantly differentially expressed, with a false discovery rate (FDR) smaller than 0.05. Out of 190, 166 of these genes were upregulated and 24 were downregulated in the *Rnf149-r^CRISPR^* condition, compared to *Tyr^CRISPR^* controls. *Rnf149-r* as well as three known pharyngeal muscle progenitor cell-specific markers, namely *Ebf*, *Htr7* and *Tbx1/10* were all significantly downregulated in the *Rnf149-r*^CRISPR^ condition (Razy-Krajka et al., 2018; Wang et al., 2019) (**Figure 4a, Supp. table 5**). By contrast, the classic cardiac determinants Nk4/Nkx2-5, Gata4/5/6 and Hand, and the heart precursor markers *Slit, Lrp4/8* and *Mmp21*, were slightly upregulated, but not significantly (**Supp. table 5**).

**Figure 4:**
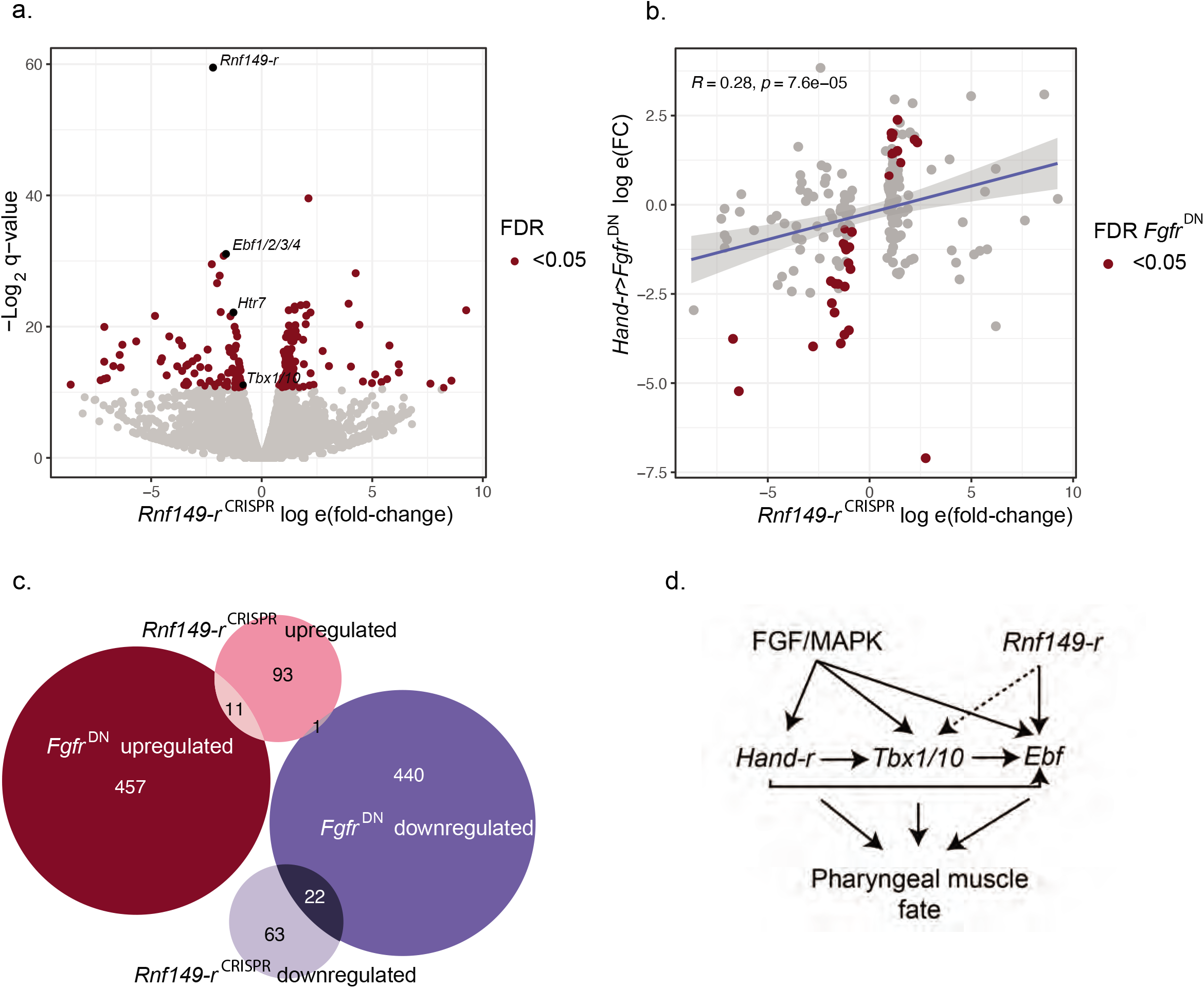
*Rnf149-r* has novel targets contributing to the cell fate choices in the cardiopharyngeal lineage. a) Volcano plot showing significantly upregulated and downregulated genes upon *Rnf149-r*^CRISPR^ compared to control (*Tyr*^CRISPR^) cardiopharyngeal lineage cells in red (FDR<0.05). Genes shown in black are well-studied ASM-specific markers (*Ebf1/2/3/4, Htr7* and *Tbx1/10*) and our target gene, *Rnf149-r*. Experiments are done in triplicates. b) Log fold-change correlations between MAPK abolishment (*Hand-r>Fgfr^DN^*) (Wang et al., 2019) and *Rnf149-r*^CRISPR^, using only differentially expressed genes in either conditions. Genes shown in red are differentially expressed in the *Rnf149-r*^CRISPR^ condition. R represents the Pearson correlation coefficient. c) Euler diagram showing overlaps of differentially expressed gene groups in *Rnf149-r*^CRISPR^ and *Hand-r>Fgfr^DN^*. Mutual enrichment p-value, calculated using Fisher’s exact test, is 4.7e-08. Odds ratio is calculated to be 3.3. d) Proposed model of *Rnf149-r*’s effect on regulation of the cardiopharyngeal lineage in *Ciona*.

As MAPK is a key regulator of fate in the cardiopharyngeal lineage, we compared the significantly changing expression levels with the overexpression of the dominant negative Fgf receptor, *Foxf>Fgfr^DN^*. We observed a positive correlation between the fold-changes for both experiments when only the significantly changing genes in the *Rnf149-r*^CRISPR^ experiment were considered (**Figure 4b**). The upregulated genes in the RNA-seq were significantly enriched in FGF-MAPK inhibited genes and the downregulated genes are significantly enriched in FGF-MAPK-dependent genes. A Fisher’s test done between the *Fgfr*^DN^ and *Rnf149-r^CRISPR^* RNA-seq show that there is significant overlap between the genes that are significantly changing in comparison to their internal controls (p-value= 4.7E-08, odds ratio = 3.3) (**Figure 4c**). These observations are consistent with a partial conversion of pharyngeal muscle progenitors to a heart-like fate in *Rnf149-r*^CRISPR^.

## Discussion

In this study, we identified *Rnf149-r* as a new regulator of cardiopharyngeal lineage development in the tunicate *Ciona*. We showed that the predicted Rnf149-r sequence contains a protein-protein interaction domain, and CRISPR/Cas9-mediated loss-of-function affects pharyngeal muscle fate specification. We developed molecular tools to study the function of this gene using CRISPR/Cas9 reagents and epistasis assays via overexpression and expression of dominant-negative reagents altering the activity of the FGF/MAPK pathway. Our analyses suggested that *Rnf149-r* acts in parallel to the FGF/MAPK pathway on shared targets. We thus uncovered a potential entry point for a novel pathway regulating cardiac vs. pharyngeal muscle fate specification.

Recent studies from our lab have shown that transcriptional inputs from FGF/MAPK signaling are required at successive stages for pharyngeal muscle specification in *Ciona* (Razy-Krajka et al., 2018; Wang et al., 2019). CRISPR/Cas9-mediated loss of *Rnf149-r* function phenocopied the loss of *Ebf* expression and pharyngeal muscle specification induced by inhibition of FGF-MAPK signaling, and an RNF149 homolog was shown to regulate Raf, we thus hypothesized that *Rnf149-r* regulates MAPK signaling. However, *Rnf149-r* loss of function did not alter expression of the multipotent progenitor marker *Hand-r*, the maintenance of which relies on continuous inputs from MEK activity. In addition, while lineage-specific bulk RNA-seq analysis of either Rnf149-r^CRISPR^ or Fgfr inhibition correlated and showed significant overlap of differentially expressed genes, including known STVC and ASMF markers such as *Htr7, Tbx1/10* and *Ebf*, there were substantial fractions of genes dysregulated by perturbation of either FGF-MAPK or Rnf149-r alone. Moreover, functional interaction assays between Rnf149-r^CRISPR^ and gain of Ras, Mek and Ets1/2 functions, indicated that Rnf149-r activity was required for each gain-of-function perturbation to cause ectopic *Ebf* expression, suggesting that *Rnf149-r* acts in parallel to FGF-MAPK-Ets, targeting a partially shared set of genes.

We note several possible future extensions of this work. First, as *Rnf149-r* is a primed heart gene, it might itself be subject to post-transcriptional regulation. Second, the role of the protein interaction domain in *Rnf149-r* is not known, and future pulldown experiments followed by Mass spectrometry-based identification of interaction partners would provide insights into Rnf149-r molecular function and the hypothesized regulatory pathway involved.

## Methods

### Ciona robusta handling

Wild *Ciona robusta* adults were obtained from M-REP (Carlsbad, CA, USA), and kept under constant light to avoid spawning. Gametes from several animals were collected separately for *in vitro* cross-fertilization followed by dechorionation and electroporation as previously described (Christiaen et al., 2009b). The embryos were cultured in filtered artificial seawater (FASW) in agarose-coated plastic Petri dishes at 18°C. We electroporated 50 μg of constructs for FACS purification (*Mesp>tagRFP, MyoD905>eGFP* and *Hand-r>tagBFP*) and 70 μg of experimental constructs (*Mesp>LacZ, Mesp>Fgfr^DN^, Mesp>Mek^S216D,S220E^*).

### CRISPR/Cas9-mediated mutagenesis

Six to eight single guide RNAs (sgRNA) per gene with Doench scores (http://crispor.tefor.net, v4.0) (Haeussler et al., 2016) higher than 60 were designed to induce mutagenesis using CRISPR/Cas9 in the B7.5 lineage as described (Gandhi et al., 2017) **(Supplementary table 4)**. The efficiency of sgRNAs was evaluated using the peakshift method as described (Gandhi et al., 2017). CRISPR/Cas9-mediated deletions were also evaluated by PCR-amplification directly from embryo lysates following electroporated with *Ef1a>nls::Cas9-Gem::nls*. sgRNAs were expressed using the *Ciona robusta U6* promoter (Stolfi et al., 2014). For each gene, two or three guide RNAs were used to total 50 μg, in combination with 25 μg of each expression plasmid. 25 μg of Mesp>nls::Cas9-Gem::nls plasmid was co-electroporated with guide RNA expression plasmids for B7.5 lineage-specific CRISPR/Cas9-mediated mutagenesis. One guide RNA was used to mutagenize *Tyrosinase* and *Neurogenin*, which are not expressed in the cardiopharyngeal lineage and thus used to control the specificity of the CRISPR/Cas9 system (Wang et al., 2019).

### Molecular cloning of *Rnf149-r^mut^ rescue construct*

Coding sequence for wild-type *Rnf149-r* (KH.C2.994) was obtained from the plasmid contained in the *C. intestinalis* full ORF Gateway-compatible clone VES66_B12. Insertion of the product into the expressing vector was performed using In-fusion (Clontech, Mountain View, CA) protocol. Oligonucleotide directed mutagenesis and two-step overlap PCRs were used to generate the point mutated form *Rnf149-r*^mut^ from the corresponding wild-type sequences. We used oligonucleotide directed mutagenesis to generate mismatches in the PAM sequences adjacent to the sgRNA targets. Because of the disturbance of a correct PAM sequence (NGG, (reverse complement CCN)), overexpressed *Rnf149-r*^mut^ is resistant to the Cas9 nuclease activity.

### Fluorescent in situ hybridization immunohistochemistry (FISH-IHC) of Ciona embryos

The following ISH probes were obtained from plasmids contained in the *C. intestinalis* full ORF Gateway-compatible clone: *Rnf149-r* (VES66_B12), *Bag3/4* (VES90_E04), *Rbm24/38* (VES87_P24), *Asb2* (VES74_P16), *Qki* (VES69_A06), *Rbms1/2/3* (VES90_E05), *Psmd14* (VES70_A18), *Ube2ql1* (VES68_H10), *Otud3* (VES70_G23), *Natn1* (VES91_K15) and: *C. intestinalis* gene collection release I: *Smurf1/2* (GC20g07), *Pcbp3* (GC07e08), *Ube2j1* (GC03l08).

PCR amplification of transcription templates was with the following oligos: M13 fw (5’-GTAAAACGACGGCCAGT-3’) and M13 rev (5’-CAGGAAACAGCTATGAC-3’). DIG- and FLUORESCEIN-labeled probes were transcribed with T7 RNA polymerase (Roche) using DIG labeling (Roche) and purified with the RNeasy Mini Kit (Qiagen). Antisense RNA probes were synthesized as described (Racioppi et al., 2014). In vitro antisense RNA synthesis was performed using T7 RNA Polymerase (Roche, Cat. No. 10881767001) and DIG RNA Labeling Mix (Roche, Cat. No. 11277073910).

Embryos were harvested and fixed at desired developmental stages for 2 h in 4% MEM-PFA (4% paraformaldehyde, 0.1 M MOPS pH 7.4, 0.5 M NaCl, 1 mM EGTA, 2 mM MgSO4, 0.05% Tween 20), rinsed in cold phosphate-buffered saline (PBS), gradually dehydrated for 1.5 h and stored in 75% ethanol at −20°C. They were then rehydrated gradually using a methanol/PBS-Tween series, and whole mount fluorescent in situ hybridization was performed as previously described (Razy-Krajka et al., 2014; Wang et al., 2013). Anti-Digoxigenin-POD Fab fragment (Roche, IN) was first used to detect the hybridized probes, then the signal was revealed using Tyramide Signal Amplification (TSA) with Fluorescein TSA Plus Evaluation Kit (Perkin Elmer, MA).

For immunohistochemistry, samples were blocked in Tris-NaCl-Blocking buffer (Blocking Reagent, PerkinElmer) for 2–4 h preceding primary antibody incubation and 1 h preceding secondary antibody incubation. Antibody solutions were prepared in Tris-NaCl-Blocking buffer and incubated for 1–2 h at room temperature, followed by an overnight incubation at 4 °C. Anti–β-galactosidase monoclonal mouse antibody (Promega, 1:500) was co-incubated with anti-mCherry polyclonal rabbit antibody (Bio Vision, Cat. No. 5993–100, 1:500) for immunodetection of *Mesp>nls::lacZ* and *Mesp>hCD4::mCherry* products respectively. Goat anti-mouse secondary antibodies coupled with AlexaFluor-555 and AlexaFluor-633 were used to detect β-galactosidase-bound mouse antibodies and mCherry-bound rabbit antibodies after the TSA reaction. Antibody washes were performed using Tris-NaCl-Tween buffer. Samples were mounted in ProLong Gold Antifade Mountant (ThermoFisher Scientific, Waltham, MA, Catalog number P36930) and stored in 4°C.

Images were acquired with an inverted Leica TCS SP8 X confocal microscope, using an HC PL APO ×63/1.30 objective. Maximum projections were processed with maximum projection tools from the Leica software LAS-AF.

### Cell dissociation and FACS-purification of Ciona robusta cells

Sample dissociation and FACS were performed as previously described (Christiaen et al., 2009a; Wang et al., 2018). Embryos and larvae were harvested at 15 hpf in 5 ml borosilicate glass tubes (Fisher Scientific, Waltham, MA. Cat.No. 14-961-26) and washed with 2 ml calcium- and magnesium-free artificial seawater (CMF-ASW: 449 mM NaCl, 33 mM Na_2_SO_4_, 9 mM KCl, 2.15 mM NaHCO_3_, 10 mM Tris-Cl pH 8.2, 2.5 mM EGTA). Embryos and larvae were dissociated in 2 ml 0.2% trypsin (w/v, Sigma, T-4799) in CMF-ASW by pipetting with glass Pasteur pipettes. The dissociation was stopped by adding 2 ml filtered ice cold 0.05% BSA CMF-ASW. Dissociated cells were passed through a 40 μm cell-strainer and collected in 5 ml polystyrene round-bottom tube (Corning Life Sciences, Oneonta, New York). Cells were collected by centrifugation at 800 g for 3 min at 4°C, followed by two washes with ice cold CMF-ASW. Cell suspensions were filtered again through a 40 μm cell-strainer and kept on ice.

Cardiopharyngeal lineage cells were labeled with *Mesp>tagRFP* and *Hand-r>tagBFP* reporters. The mesenchyme cells were counter-selected using *MyoD905>GFP*. Dissociated cells were loaded in a BD FACS AriaTM cell sorter. 488 nm laser, FITC filter was used for GFP; 407 nm laser, 561 nm laser, DsRed filter was used for tagRFP and Pacific BlueTM filter was used for tagBFP.

### RNA-seq library preparation, sequencing and analysis

To profile transcriptomes of FACS-purified cells from *Rnf149-r^CRISPR^* and control samples, 1,000 cells were directly sorted in 100 μl lysis buffer from the RNAqueous-Micro Total RNA Isolation Kit (Ambion). For each condition, samples were obtained in three biological replicates. The total RNA extraction was performed following the manufacturer’s instruction. The quality and quantity of total RNA was checked using Agilent RNA ScreenTape (Agilent) using the 4200 TapeStation system. RNA samples with RNA Integrity Number (RIN) > 8 were kept for downstream cDNA synthesis. 250–2000 pg of total RNA was loaded as a template for cDNA synthesis using the SMART-Seq v4 Ultra Low Input RNA Kit (Clontech) with template switching technology. RNA-Seq Libraries were prepared and barcoded using Ovation Ultralow System V2 (NuGen). Six barcoded samples were pooled in one lane of the flow cell and sequenced by Illumina NextSeq 750 (MidOutput run). Paired-end 75 bp length reads were obtained from all the bulk RNA-seq libraries. Bulk RNA-seq libraries were aligned using STAR 2.7.0a (Zhang et al., 2018) with the parameters ‘--runThreadN 6 --outSAMtype BAM SortedByCoordinate \ --outSAMunmapped Within \ --outSAMattributes Standard’. Counts were obtained using featureCounts, a function of subread (Liao et al., 2014; Liao et al., 2019). Differential expression was calculated using DESeq2 (Love et al., 2014).

## Data availability

The RNA-seq data were deposited in the Gene Expression Omnibus (GEO) under accession GSE171152. Previously published bulk RNA-seq data that was used here for comparison in **Figure 4** can be found on GEO with accession GSE99846.

## Acknowledgements

B.V. acknowledges funding by American Heart Association grant #18PRE33990254, NYU Fleur Strand Graduate Fellowship and NYU GSAS Dean’s Dissertation Fellowship. C.V. acknowledges funding by the National Institutes of Health 5R35GM127089. L.C. acknowledges funding by the Leducq Foundation, award 15CVD01, and by the National Institute of Health, award R01 HL108643. We thank Yelena Bernadskaya for the critical reading of and feedback on the manuscript. We thank Alberto Stolfi and Shashank Gandhi for *Nova*^CRISPR^ and *Nova* riboprobe and Manar Elzaky for *Rnf149-r* structure analysis. We thank NYU Genomics Core facility for technical support for FACS.

## Supplemental Material

**Supplementary Table 1:**
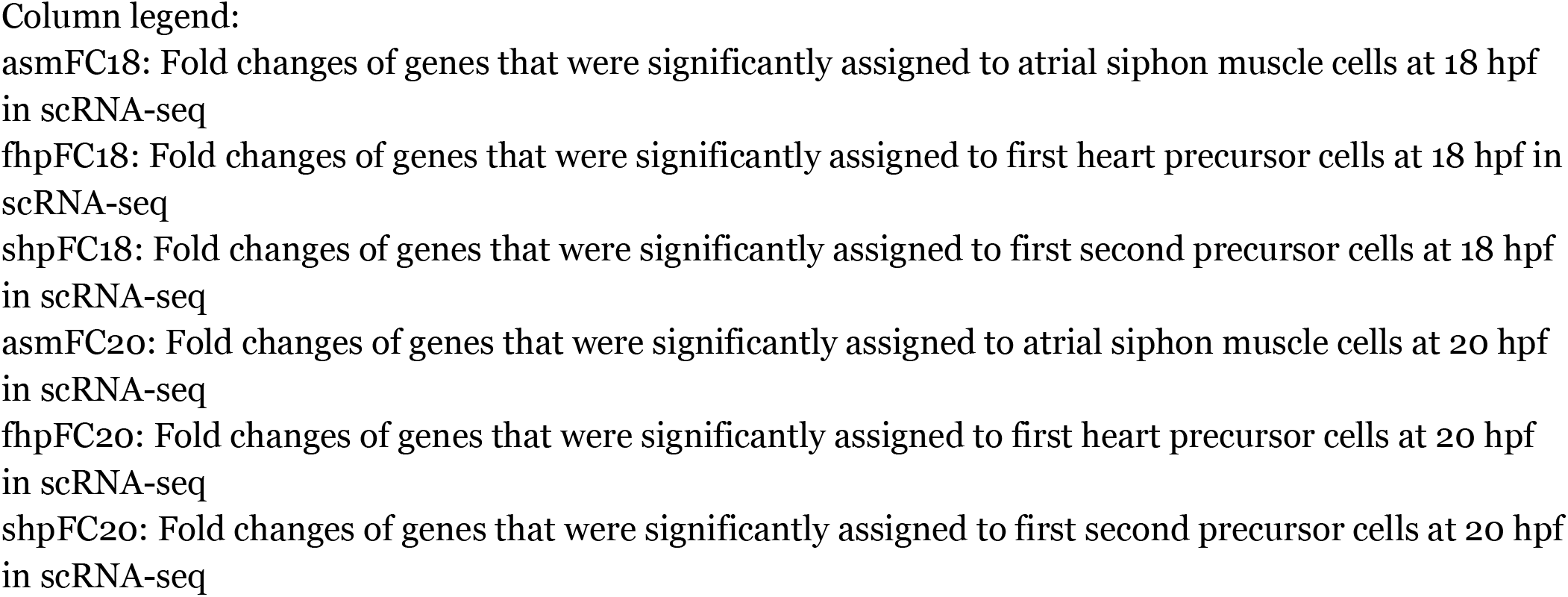
Supp table 1.xlsx

**Supplementary Table 2:** Supp Table 2

| Hours post fertilization (hpf) | FABA Stage (Hotta et al., 2007; Hotta et al., 2020) | Developmental Stage |
| --- | --- | --- |
| 12 | Stage 23 | late tailbud I |
| 15 | Stage 24/25 | late tailbud II-III |
| 18 | Stage 26/27 | hatching larva |
| 20 | Stage 28 | early swimming larva |
| 26 | Stage 30 | early swimming larva |

**Supplementary Table 3:**
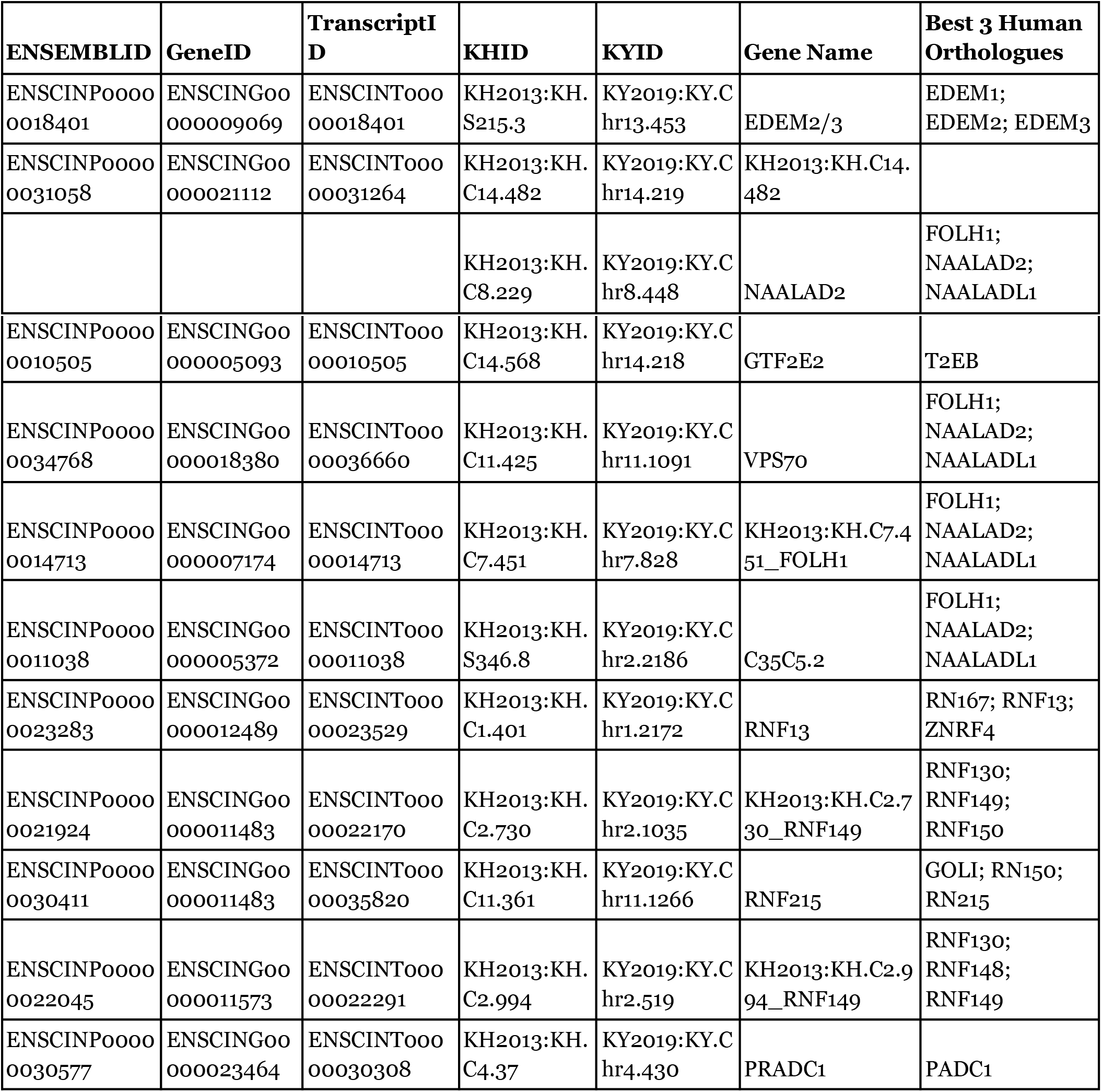
Supp Table 3 Comprehensive list of *Ciona* proteins that have recognized PA-domains. ENSCINP00000031058 maps to 2 KH IDs.

**Supplementary Table 4:**
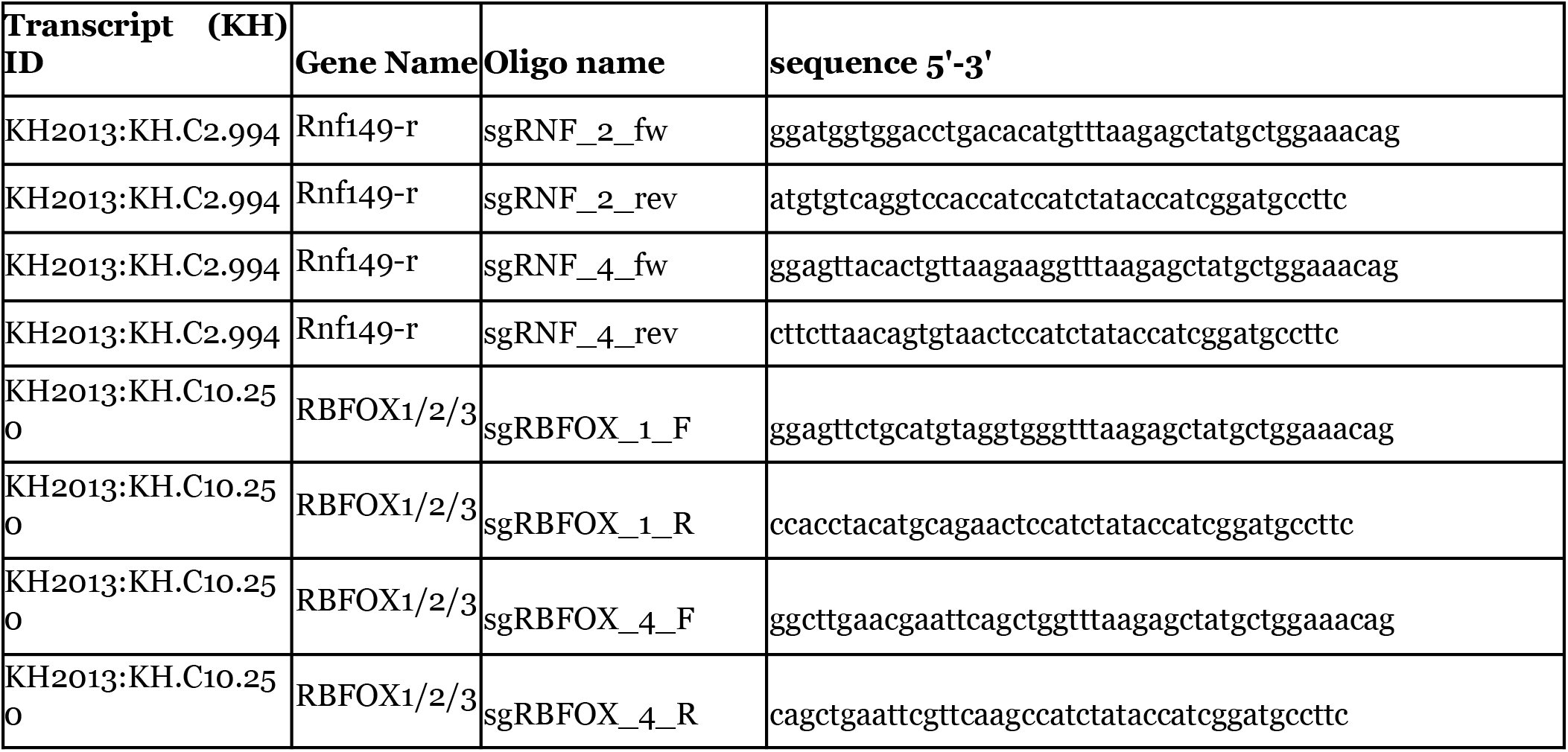
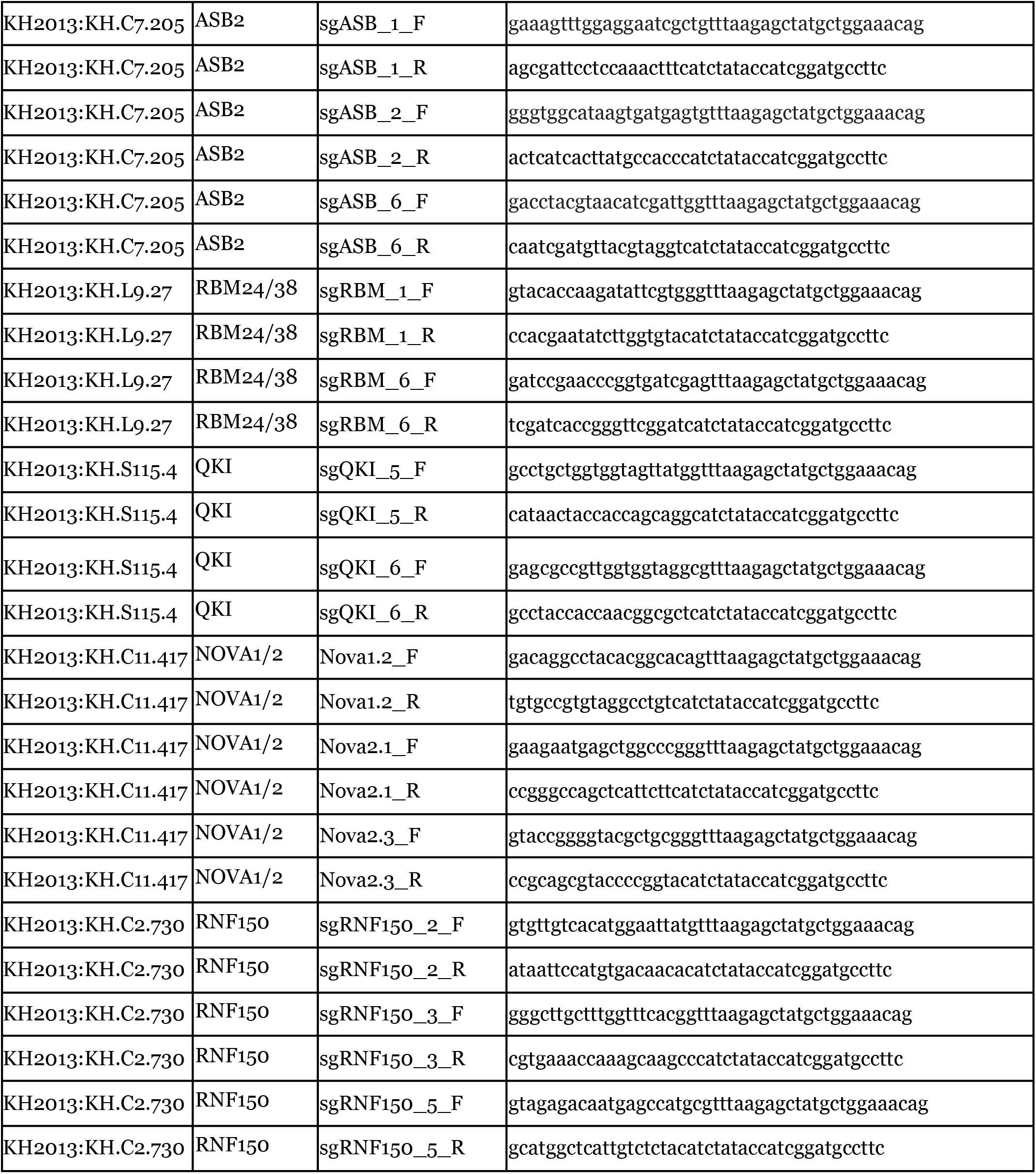
Genes tested and primers used to construct sgRNAs to validate post-transcriptional regulators detected by scRNA-seq.

**Supplementary Table 5:**
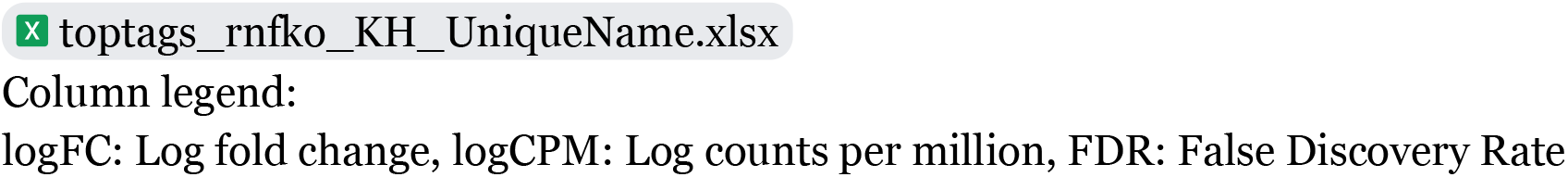
*Tyr*^CRISPR^ vs. *Rnf149-r*^CRISPR^ RNA-seq gene list with log fold-change (logFC), p-value (PValue) and false discovery rate (FDR) values.

## Supplemental figure legends

**Figure S1:**
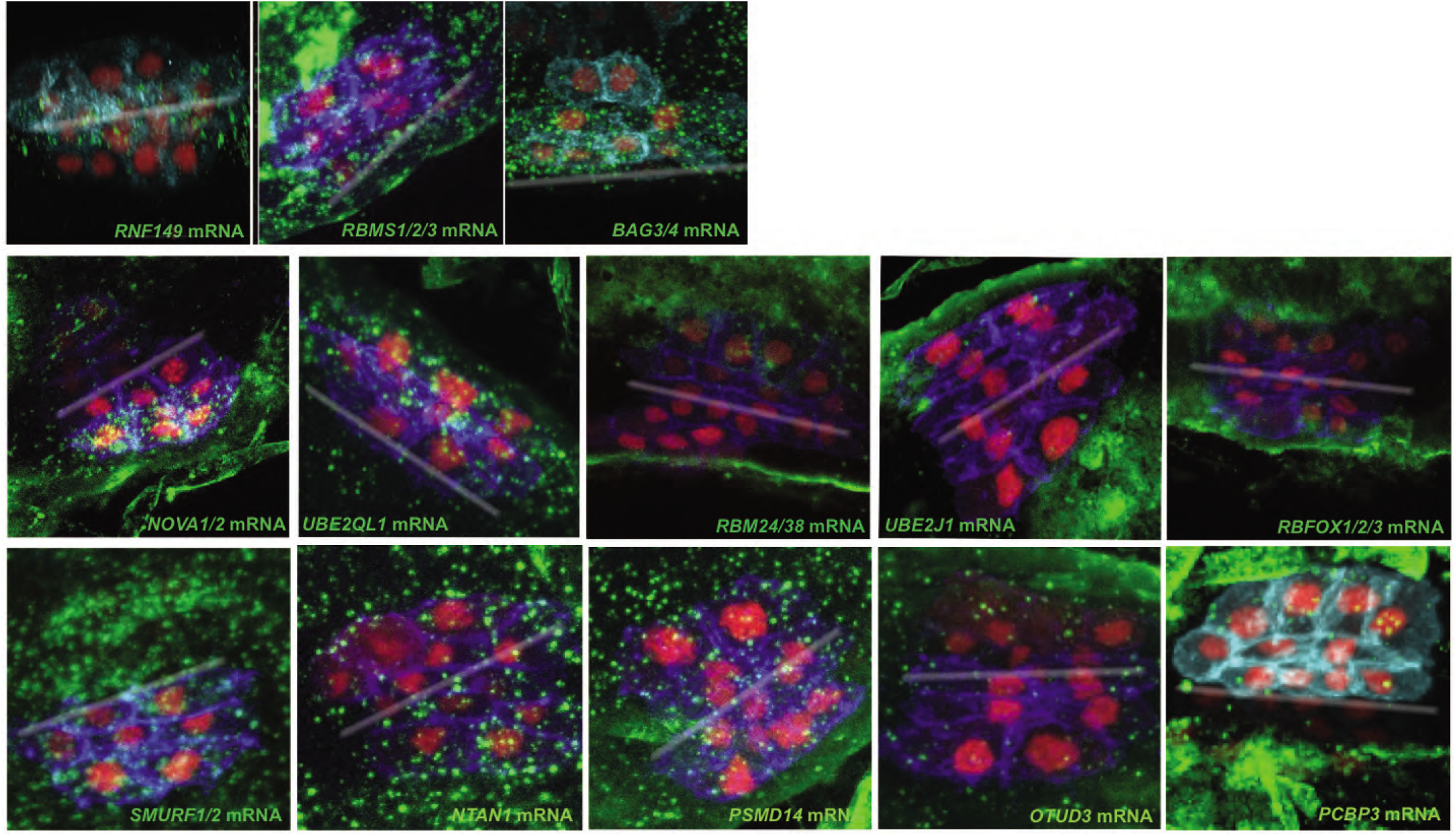
FISH validation of the expression for candidate genes. First row is predicted heart genes, second and third row predicted ASM genes. mRNAs are visualized by whole mount fluorescent in situ hybridization (green). Nuclei of cells are marked by Mesp>nls::LacZ and revealed by anti beta-galactosidase antibody (red). Mesp>hCD4::mCherry accumulates in the cell membrane and is revealed by anti mCherry antibody (blue). White line indicates the midline.

**Figure S2:**
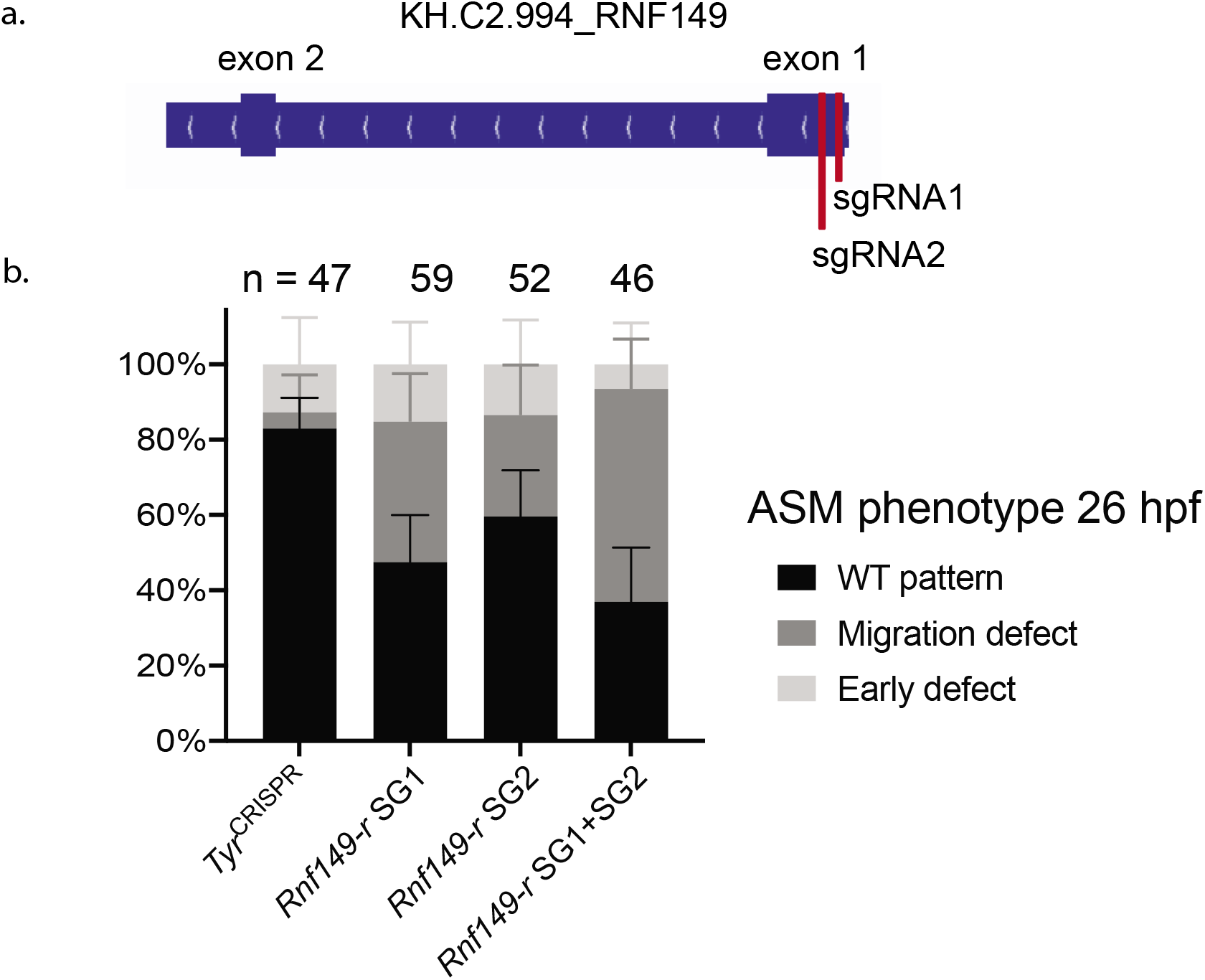
FISH validation and violin plots of the expression for *Rnf149* at different developmental time points. a) mRNAs are visualized by whole mount fluorescent in situ hybridization (green). Cells are marked by *Mesp>mCherry*. b) Violin plots represent the distribution of the expression of indicated genes in defined cell clusters (Wang et al., 2019). Each cell is represented by a black dot. On the x-axis, the numbers indicate hours post fertilization (hpf). Scale bar, 50 μm.

**Figure S3:**
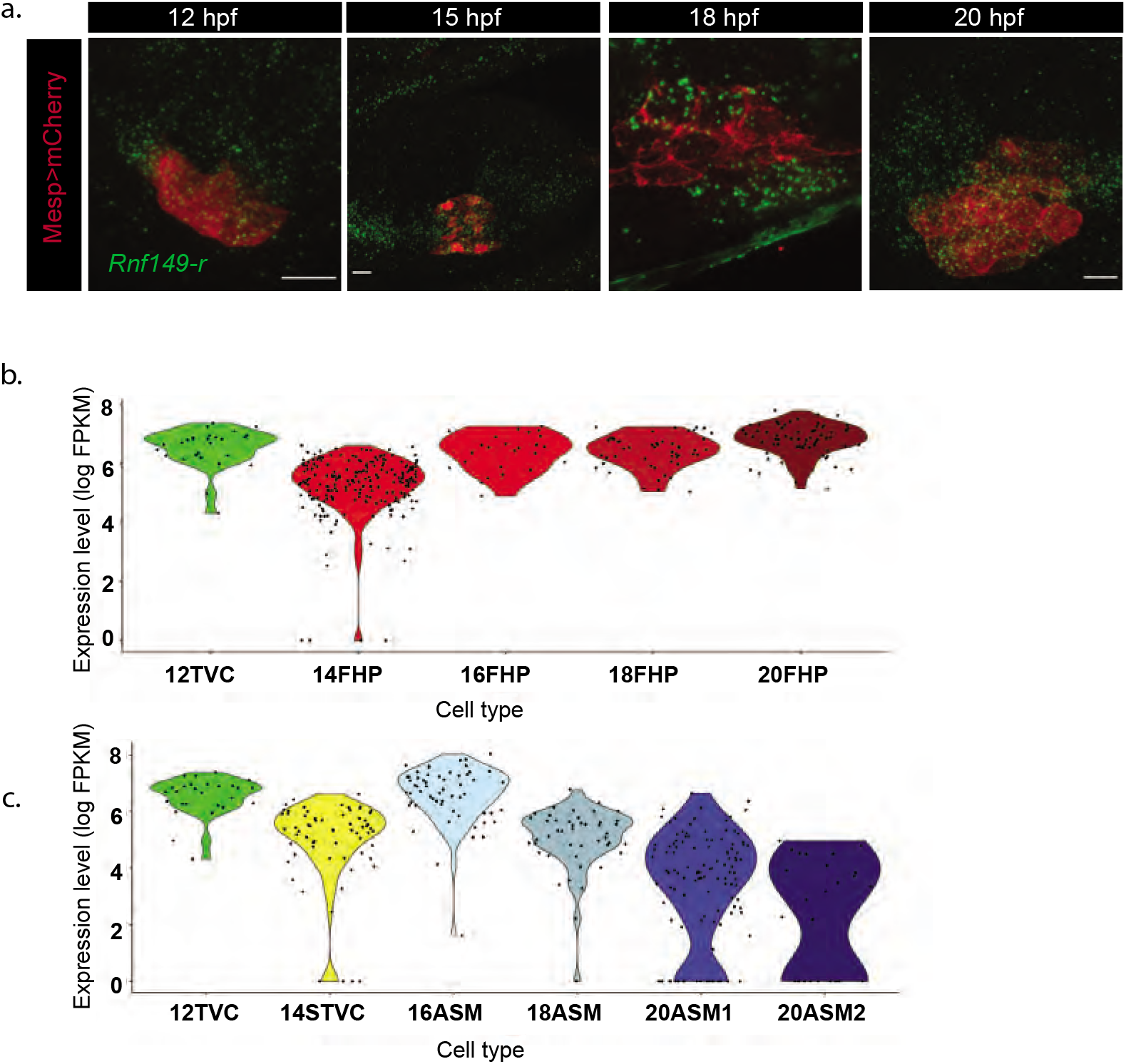
Both sgRNAs used to generate *Rnf149-r*^CRISPR^ contribute to the phenotype. a) Schematic of the *Rnf149-r* gene and the positions of the two sgRNAs used. b) Histogram with phenotype proportions. “n=” represents the total numbers of individual halves scored per condition. Error bars represent 95% Wilson method of confidence interval for proportions.

**Figure S4:**
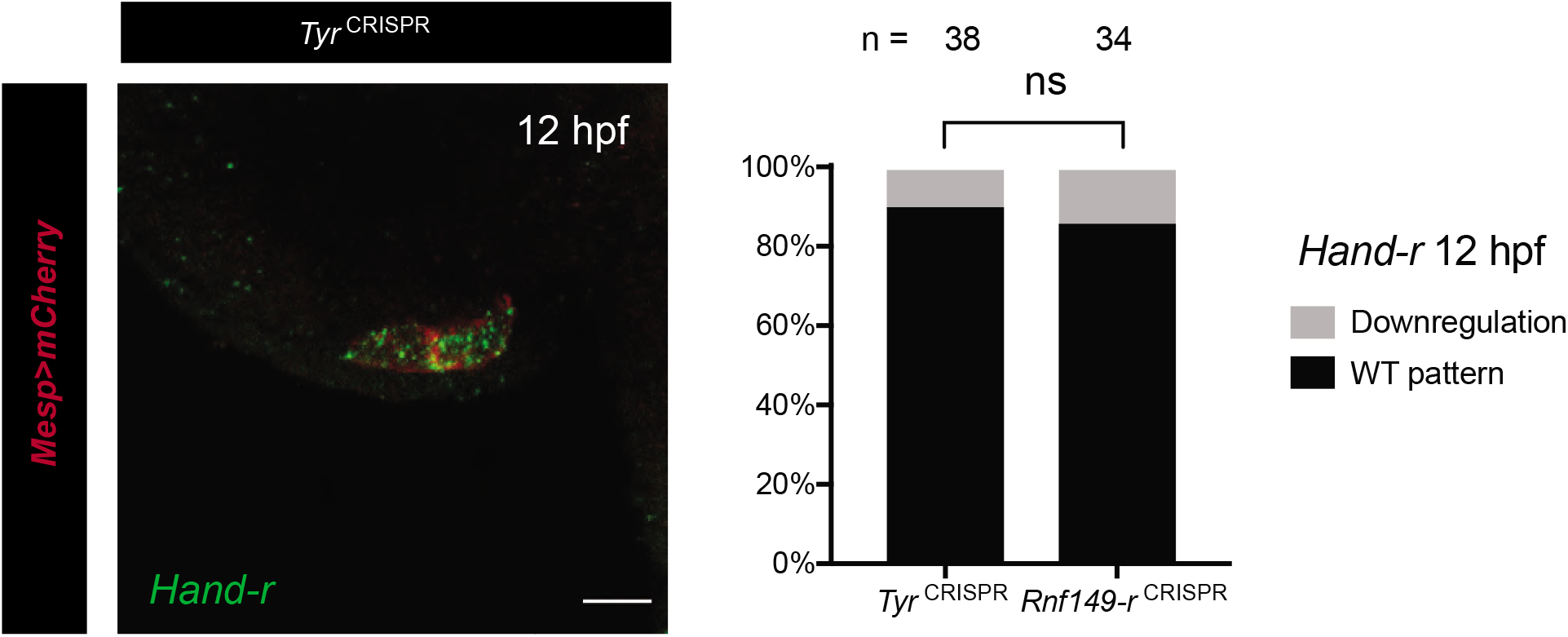
*Rnf149-r*^CRISPR^ does not alter the TVC marker *Hand-r* expression at 12 hpf. Left panel shows the in situ hybridization against a *Hand-r* probe in green. *Tyr*^CRISPR^ used as control. Cardiopharyngeal lineage marked by mCherry driven by *Mesp* and revealed by mCherry antibody in red. H2B::mCherry accumulates in the nuclei. Scale bar, 50 μm. Right panel shows the corresponding histogram with phenotype proportions. “n=” represents the total numbers of individual halves scored per condition.

**Figure S5:**
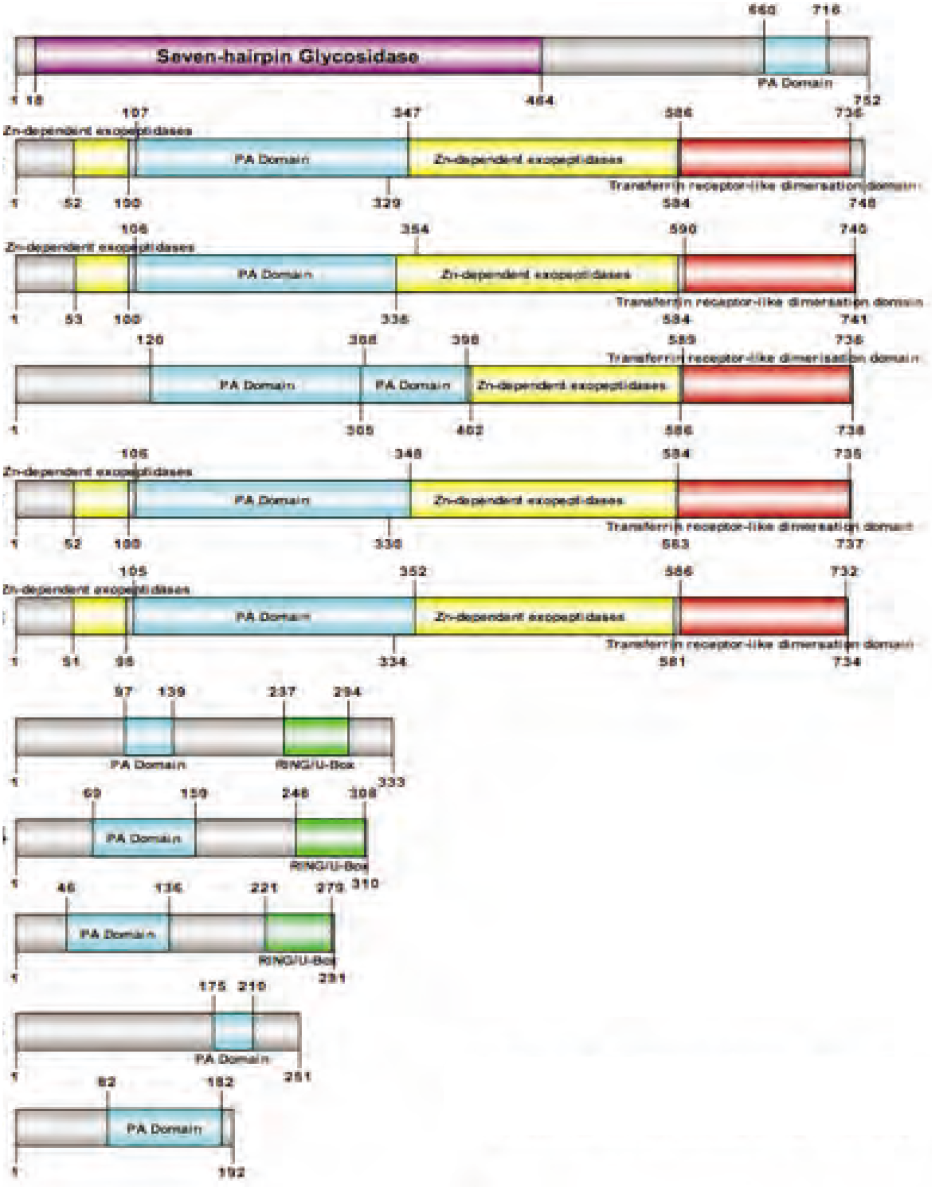
There are 11 PA domain-containing genes in the Ciona genome. PA domains shown in light blue, RING domains shown in green, Zn-independent exopeptidase domains shown in yellow, transferrin receptor-like dimerization domains shown in red and seven-hairpin glycosidase domain shown in purple.

**Figure S6:**
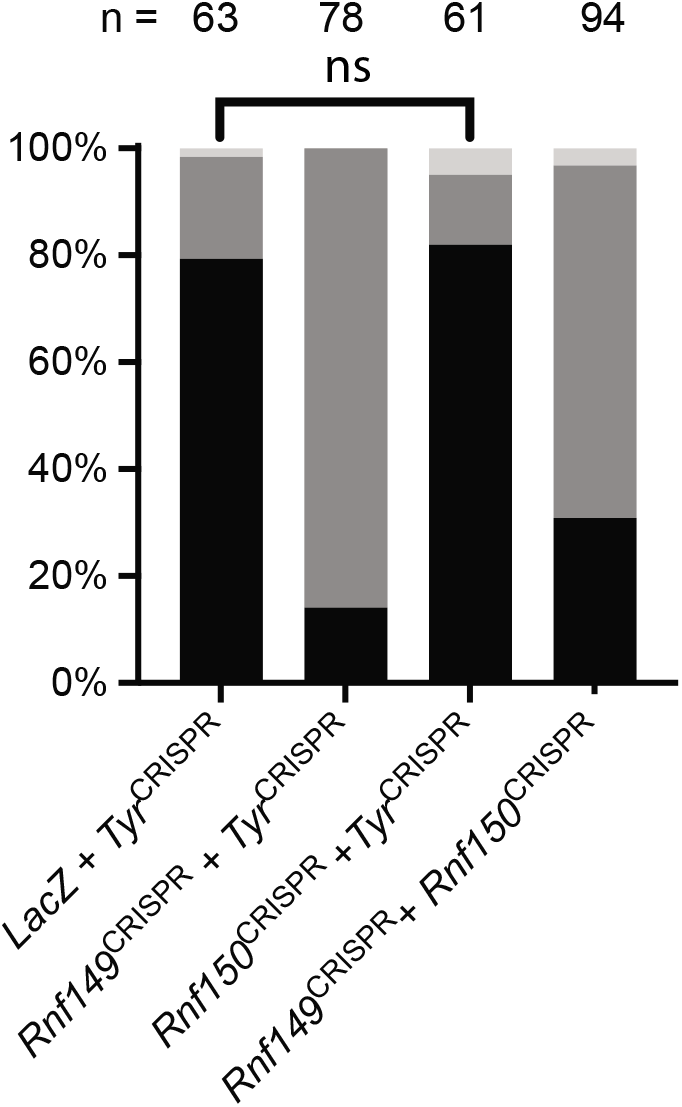
*Rnf150*^CRISPR^ did not result in significant phenotypic deformities at 26 hpf. Histogram with phenotype proportions. “n=” represents the total numbers of individual halves scored per condition.

**Figure S7:**
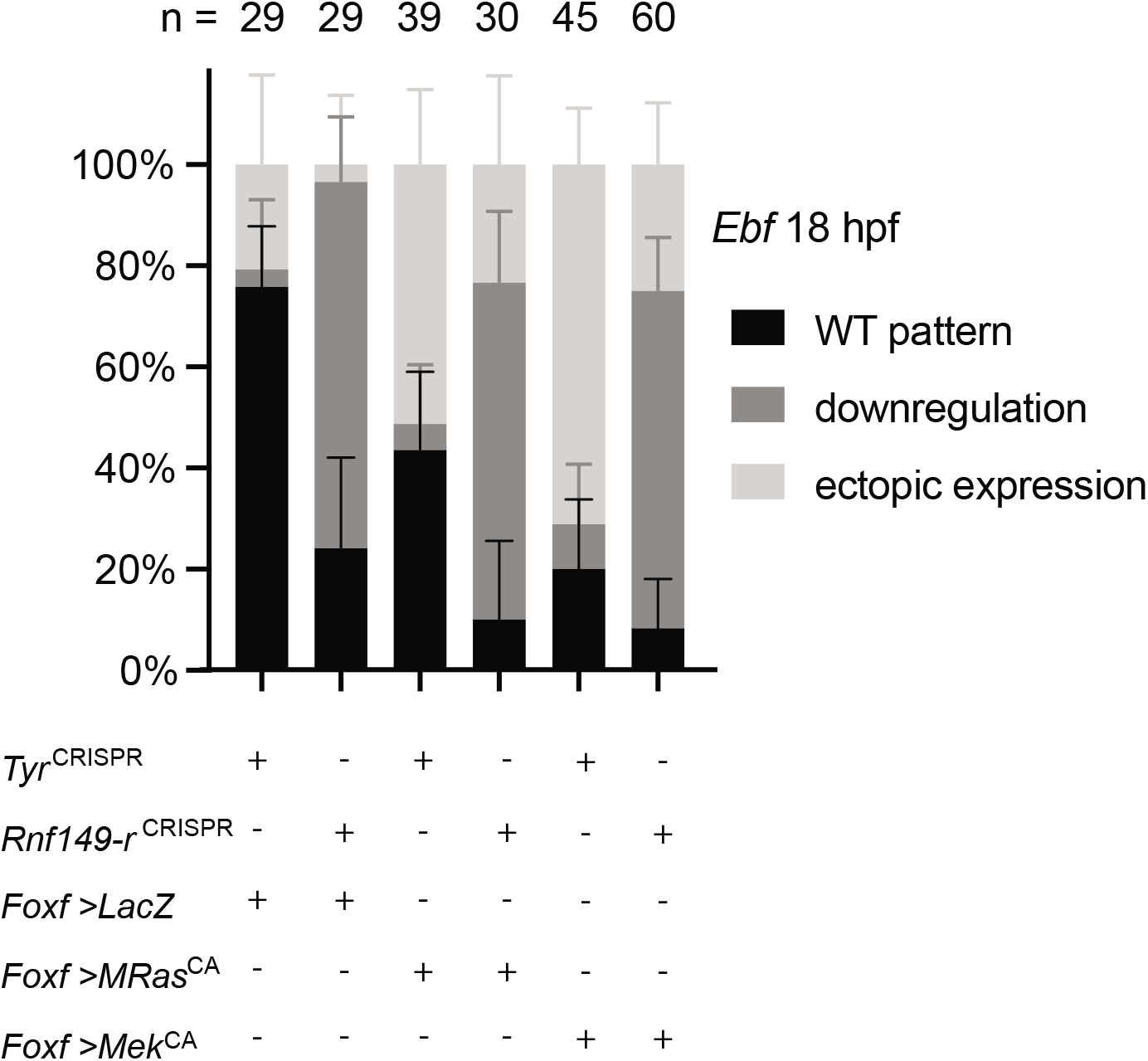
The histogram for the TVC-specific *FoxF* enhancer labeling, FoxF>LacZ, with phenotype proportions. “n=” represents the total numbers of individual halves scored per condition. Error bars represent 95% Wilson method of confidence interval for proportions.

